# Predicting the N400 ERP component using the Sentence Gestalt model trained on a large scale corpus

**DOI:** 10.1101/2021.05.12.443787

**Authors:** Alessandro Lopopolo, Milena Rabovsky

## Abstract

The N400 component of the event related brain potential is widely used to investigate language and meaning processing. However, despite much research the component’s functional basis remains actively debated. Recent work showed that the update of the predictive representation of sentence meaning (semantic update, or SU) generated by the Sentence Gestalt model (McClelland, St. John, & Taraban, 1989) consistently displayed a similar pattern to the N400 amplitude in a series of conditions known to modulate this event-related potential. These results led Rabovsky, Hansen, and McClelland (2018) to suggest that the N400 might reflect change in a probabilistic representation of meaning corresponding to an implicit semantic prediction error. However, a limitation of this work is that the model was trained on a small artificial training corpus and thus could not be presented with the same naturalistic stimuli presented in empirical experiments. In the present study, we overcome this limitation and directly model the amplitude of the N400 elicited during naturalistic sentence processing by using as predictor the SU generated by a Sentence Gestalt model trained on a large corpus of texts. The results reported in this paper corroborate the hypothesis that the N400 component reflects the change in a probabilistic representation of meaning after every word presentation. Further analyses demonstrate that the SU of the Sentence Gestalt model and the amplitude of the N400 are influenced similarly by the stochastic and positional properties of the linguistic input.

## Introduction

The N400 event-related potential (ERP) component is a negative deflection at centro-parietal electrode sites peaking around 400 ms after the onset of a word or another potentially meaningful stimulus. Its amplitude has been shown to be affected by a wide variety of linguistic variables. For instance, N400 amplitudes tend to decrease over the course of a sentence (van Petten & Kutas, 1990). Smaller amplitudes are observed for targets after semantically similar or related as compared to unrelated primes, and for repeated words as compared to a first presentation (Bentin, McCarthy, & Wood, 1985). The N400 shows larger amplitude to congruent continuations with lower as compared to higher cloze probability (Kutas & Hillyard, 1984). In general, the amplitude of the N400 seems to be sensitive to the stochastic properties of the word, both in terms of lexical frequency and surprisal (van Petten & Kutas, 1990; Parviz, Johnson, Johnson, & Brock, 2011; Frank, Otten, Galli, & Vigliocco, 2015), with lower amplitudes observed for high frequency words and for words with lower surprisal. Despite the large body of data on N400 amplitude modulations and the agreement that the N400 is related to meaning processing, the computational principles and processing mechanisms underlying N400 amplitude generation are as yet unclear. Various theories propose, e.g., that the N400 reflects, among others, lexical-semantic access or semantic integration processes (Kutas & Federmeier, 2011).

Recent studies linking the N400 to computational models in order to better understand the underlying mechanisms can be ascribed to two broadly defined categories, (neuro)cognitively motivated small scale models linking the N400 to internal processes in the models and large-scale natural language processing (NLP) based models computing surprisal. As examples of the latter type, Frank et al. (2015); Aurnhammer and Frank (2019); Merkx and Frank (2020) have shown that the N400 amplitude is significantly influenced by word-level surprisal, computed by a variety of language model implementations trained on a next-word prediction task. An advantage of these models is that they are trained on large linguistic datasets, approximating human language exposure so that they can be directly linked to empirical N400 data. However, the measure that is used to predict N400 amplitudes, surprisal, is an output measure of the models that can be computed in many different ways and thus does not directly speak to the internal cognitive processes and neural activation dynamics underlying N400 amplitudes, which is our main interest here.

On the other hand, cognitively motivated computational models of sentence comprehension that have been used to model the N400 (Brouwer, Crocker, Venhuizen, & Hoeks, 2017; Brouwer, Delogu, Venhuizen, & Crocker, 2021; Fitz & Chang, 2019; Rabovsky et al., 2018; Rabovsky & McClelland, 2020; Rabovsky, 2020) link N400 amplitudes to internal hidden layer activation processes and dynamics, thereby providing computationally explicit (neuro)cognitive theories of the processes giving rise to the N400. Most of these models were trained to map sequentially incoming words to estimated sentence meaning, which is arguably an important part of human language comprehension. On the downside, these cognitively motivated models proposed to account for N400 amplitudes have as of yet been only trained on small artificial language corpora, which makes the relationship to empirical N400 data somewhat abstract.

Here, we aim to combine the advantages of both approaches by training a cognitively motivated model of language processing that has been used to model the N400, the Sentence Gestalt (SG) model (Rabovsky et al., 2018; Rabovsky & McClelland, 2020; Rabovsky, 2020), on a large-scale language corpus, to directly test our computationally explicit (neuro)cognitive theory of the processes underlying N400 amplitudes on single trial empirical ERP data.

Specifically, Rabovsky et al. (2018) proposed an explanation of the N400 ERP component in terms of update of a probabilistic representation of meaning as captured by the change of the inner states of the SG model, a connectionist model of language processing that maps a sentence to its corresponding event (McClelland et al., 1989). At every given moment during sentence processing, this representation not only contains information provided by the words presented so far, but also an approximation of all features of the sentence meaning based on the statistical regularities in the model’s environment internalized in its connection weights. Rabovsky et al. (2018) showed that the SG model update simulates a number of N400 effects obtained in empirical research including the influences of semantic congruity, cloze probability, word position in the sentence, reversal anomalies, semantic and associative priming, categorically related incongruities, lexical frequency, repetition, and interactions between repetition and semantic congruity. These results foster the idea that N400 amplitudes reflect surprise at the level of meaning, defined as the change in the probability distribution over semantic features in an integrated representation of meaning occasioned by the arrival of each successive constituent of a sentence.

In the present study, we attempt to directly predict the amplitude of the N400 generated during sentence processing by using as predictor the update of the inner representation of a SG model trained on a large corpus of naturalistic texts. We used EEG data collected while subjects were asked to read sentences extracted from narrative texts. Such stimuli were not designed to elicit specific conditions, but to be close to everyday conditions faced by humans.

The hypothesis that N400 amplitudes reflect the change in a probabilistic representation of meaning after every word presentation is supported by the results reported in the present study, which show that the SG model update successfully predicts N400 amplitudes. Moreover, in order to further investigate the relations between the inner dynamics of the SG model and the N400, we conducted analyses aimed at assessing whether the update of the inner representation generated by the model and the amplitude of the N400 component are influenced in the same way by the word frequency, surprisal and position of the words making up the stimulus.

## The Sentence Gestalt Model

As a model of language processing, the SG model maps sentences to a representation of the described event approximated by a list of role-filler pairs representing the action, the various participants (e.g., agent and patient) as well as information concerning, for instance, the time, location, and the manner of the event described by the sentence itself (McClelland et al., 1989).

1. The boy ate soup during lunch *agent action patient time*

For instance, when processing Sentence 1, the SG model recognizes that *ate* is the action, and that *the boy* and *soup* are its agent and patient respectively and that *at lunch* is a modifier specifying the moment when the event takes place.

The SG model consists of two components: an update network (encoder) and a query network (decoder), as described in Fig. 1. The update network sequentially processes each incoming word to update activation of the SG layer, which represents the meaning of the sentence after the presentation of each word as a function of its previous activation and the activation induced by the new incoming word. The query network, instead, extracts information concerning the event described by the sentence from the activation of the SG layer. The sentence comprehension mechanism is implemented in the update network. The query network is primarily used for training.

**Fig. 1:**
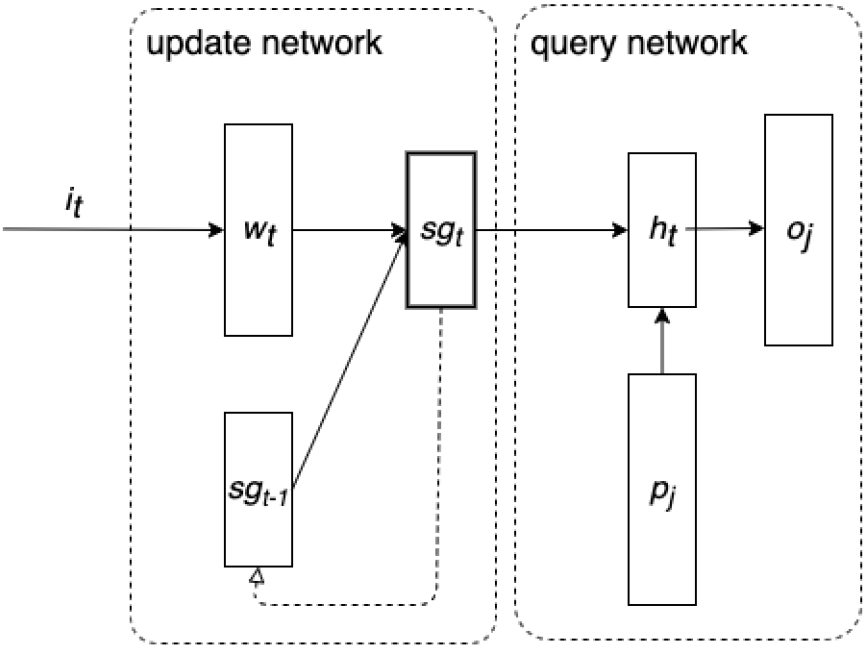
The architecture of the Sentence Gestalt Model, with the **update network** on the left hand-side and the **query network** on the right hand-side.

In the present study, the **update network** of the SG model is composed of an input layer, which generates a vectorial representation 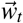 for each input word *i_t_* of the incoming sentence, and a recurrent layer implemented as a long short-term memory (LSTM) unit generating a SG representation 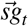 as a function of 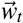 and previous gestalt 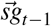 (Hochreiter & Schmidhuber, 1997). LSTM have the advantage of being better at processing long and complex sentences compared to traditional recurrent layers, and being still simpler in structure and number of parameters compared to even more performative types of deep learning components (e.g. Transformers). In the original formulation of the update network, the SG representation 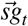 is obtained from the activity of a hidden layer which combines the previous SG representation 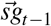 and the vector of the current word 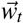. The adoption of LSTM in the present implementation of the SG model is justified by the significantly more complex nature of the training material as compared to the original implementation of the model which was trained on short and artificially generated sentences. Nonetheless, the basic principles of the update network are retained in this novel formulation, since in both cases the network is essentially a recurrent neural network encoding a string of words as a function of the current presented word and the words preceding it. The **query network** instead has the same structure as the original SG model implementation. Its hidden state 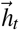 is generated by combining the SG vector 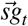 and probe vectors 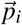. The output 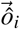 of the query network is generated from the hidden state 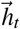.

The **task** the SG model is asked to perform is to map a sentence to its corresponding situation or event, defined as a list of role-filler pairs representing an action or state, its participants (e.g. agent, patient, recipient), and eventual modifiers. A sentence is defined as a sequence of words, each represented as an integer *i_t_*, defined on a vocabulary associating a unique index to every word. Figure 2 exemplifies the mapping from words to event performed, word-by-word, by the SG model.

**Fig. 2:**
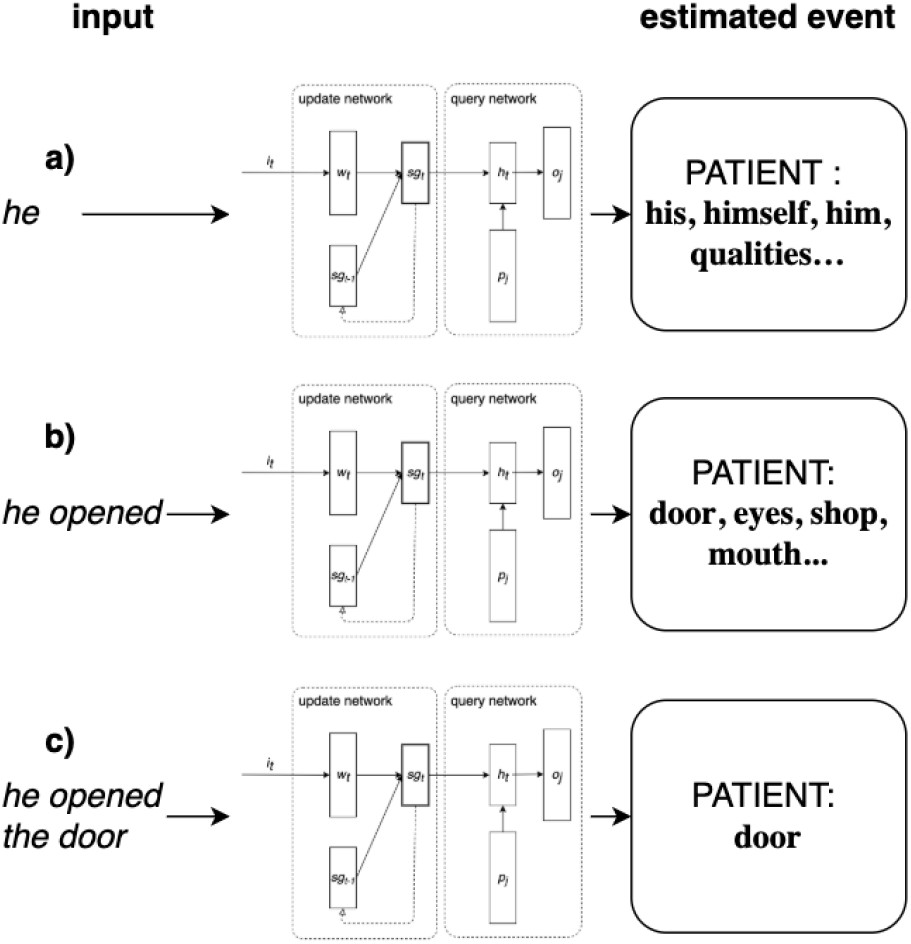
Example of how the SG model processes the sentence *he opened the door*. The model is trained on identifying the roles and fillers constituting the event described in the sentence. Here, for reasons of space, we focus only on mapping filler to role *patient*, but the model also estimates the agent and action of the event. Sentences are presented word-by-word. Initially (a), the model is first presented only with word *he*, leading to a list of wrong predicted patient fillers. Subsequently (b), the model is presented with word *opened*, causing the prediction of potentially correct fillers. Finally (c), when the model is presented with the whole sentence, it converges on the correct patient of the described event: *door*.

As shown in Fig. 3.a, the event consists of a set of rolefiller vectors 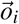, each of which consists of the concatenation of the feature representation of a word and a one-hot vector of the role of that word in the context of the event described by the sentence. Sentence 1 above consists of a sequence of 7 one-hot word representation vectors. Its event contains 4 role-filler combinations representing each role of its event (agent, action, patient, time) with its corresponding concept filler (boy, eat, soup, lunch).

**Fig. 3:**
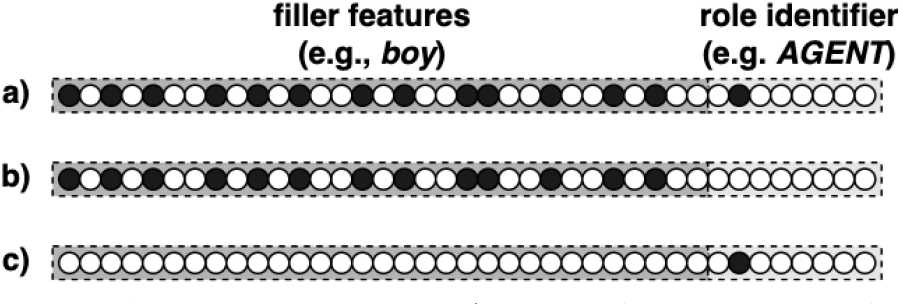
The role-filler vector 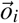 (a), and its corresponding two types of probes 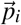 (b) and (c). The left hand-side of the vectors correspond to the embedding representation of the filler concept, whereas the right hand-side to the one-hot representation of the thematic role played by the filler. When probing for the thematic role, probe (b) is presented. When probing for the filler instead, probe (c) is presented. In both cases the SG model is expected to produce the full role-filler vector (a).

During **training**, the model is presented with sentences (fed word by word to the input layer) such as ‘The boy ate soup during lunch’. Every time a word is presented, the model is probed concerning the event described by the sentence. The model is probed concerning the complete event, even if the relevant information has not yet been presented at the input layer. A probe consists of a vector 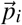 of the same size of a corresponding role-filler vector 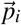, but with either the thematic role identifier zeroed (Fig. 3.b) – if probing for roles –, or filler features zeroed (Fig. 3.c) – if instead probing for fillers. Responding to a probe consists therefore of completing the role-filler vector. When probed with either a thematic role (e.g., agent, action, patient, location, or situation; each represented by an individual unit at the probe and output layer) or a filler, the model is expected to output the complete role-filler vector. Fillers are represented using word embeddings obtained by binarizing *Fasttext*, a computational semantic model representing 1 million words and trained on both the English Wikipedia and the Gigaword 5 corpora (Bojanowski, Grave, Joulin, & Mikolov, 2017). The discrepancies between the observed role-filler vector 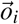, and generated output 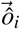 is computed using cross-entropy and is back-propagated through the entire network to adjust its parameters in order to minimize the difference between model-generated and correct output. Binarization of the filler semantic feature representations was performed in order to allow for a probabilistic interpretation of the model generated activation of semantic feature units afforded by the cross-entropy error used during training.

**Fig. 4:**
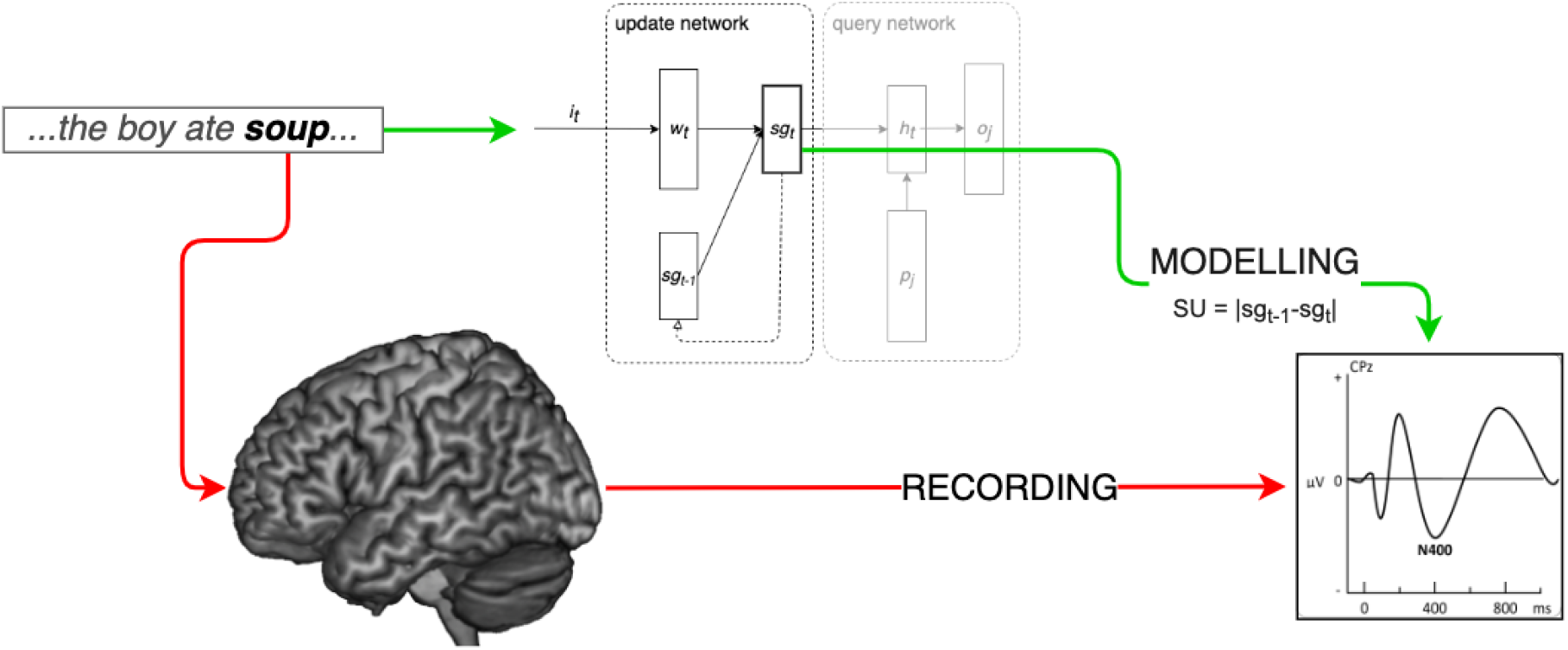
This study aims to model the amplitude of the N400 as a function of the update of a probabilistic semantic representation (SU) generated by a SG model trained on a large scale corpus of naturalistic texts. N400 amplitudes elicited by reading a sentence (e.g., *the boy ate soup*…) are matched by the SU generated by the model.

## Materials and methods

Previous implementations of the SG model have been trained on artificially generated sentences, constructed in order to sample basic word and role-filler combinations. In the present study, the model is trained on a large-scale training corpus approximating the real life language experience of human participants. This approach allows for simulation of empirical experiments with the exact same stimuli on a single-trial basis.

### Training corpus and hyper-parameters

The SG model was trained on the British National Corpus section of the Rollenwechsel-English (RW-eng) corpus (Sayeed, Shkadzko, & Demberg, 2018). The RW-eng corpus is annotated with semantic role information based on PropBank roles (Palmer, Gildea, & Kingsbury, 2005) and obtained from the output of the SENNA semantic role labeller (Collobert et al., 2011; Collobert, 2011) and the MALT syntactic parser (Nivre, 2003). Each sentence annotation consists of the list of event frames it describes. An event frame is defined by its main predicate (usually a finite verb) and its arguments. Following PropBank, RW-eng frames can contain arguments of 26 types spanning from agent, patient, benefactive, starting and end point and a series of modifiers describing the temporal, locational, causal, final and modal circumstances of the event. Therefore, the SG model in this study is trained on mapping each RW-eng sentence to its PropBankstyle event structure as provided in the RW-eng corpus. For more detail on the argument structure proposed by PropBank we refer to Palmer et al. (2005). A sentence can contain multiple event frames.

The parameters of the SG model were optimized using Adamax (Kingma & Ba, 2015) with learning rate equal to 0.0005. The data was split in mini-batches of 32 sentences each. Training was conducted for a maximum of 400 epochs on 90% of the batches, the remaining 10% was kept for validation. Only sentences having between 4 and 25 words and having a maximum of 10 frames were used for training. Sentence length and number of frames were constrained in order to limit the number of complex subordinate events and to facilitate the mini-batch training. That yielded a total of 64017 training batches (2048544 sentences) and 7113 validation batches (227616 sentences).

The size of the hidden layers (including the SG layer) was 2400, whereas the input layer generates per-word embeddings of size 600 for the 8000 word forms accepted. The probe and output layers had size 337 due to the concatenation of the 300-size binarized embedding vector, the frame number and the argument type.

### EEG dataset

The elecrophysiological recordings of the N400 were obtained from an EEG dataset provided by Frank et al. (2015). The dataset consists of data collected from twenty-four participants (10 female, mean age 28.0 years, all right handed and native speakers of English) while they were reading sentences extracted from English narrative texts.

The stimuli consisted of 205 sentences (1931 word tokens) from the UCL corpus of reading times (Frank, Monsalve, Thompson, & Vigliocco, 2013), and originally from three little known novels. The sentences were presented in random order, word by word. The N400 amplitude for each subject and word token was defined as the average scalp potential over a 300-500 ms time window after word onset at electrode sites in a centro-parietal region of interest.

For further details regarding the stimuli see Frank et al. (2013). More detailed information regarding the EEG dataset, its stimulation paradigm and preprocessing can instead be found in Frank et al. (2015).

## Analyses

In order to assess the hypothesis that the N400 component reflects the change of the neural representation of predicted meaning after every word presentation, we fitted a linear mixed effect model with the aim of predicting the amplitude of the N400 as a function of the update of the semantic representation generated by the SG model during language processing (the Semantic Update or SU). The SU is computed as the mean absolute error between 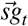 and 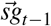 generated by the SG layer of the update network. Fig. 5 presents a graphical summary of our approach. Since the N400 is a negative deflection of the electrophysiological signal, the SU is multiplied by – 1.

**Fig. 5:**
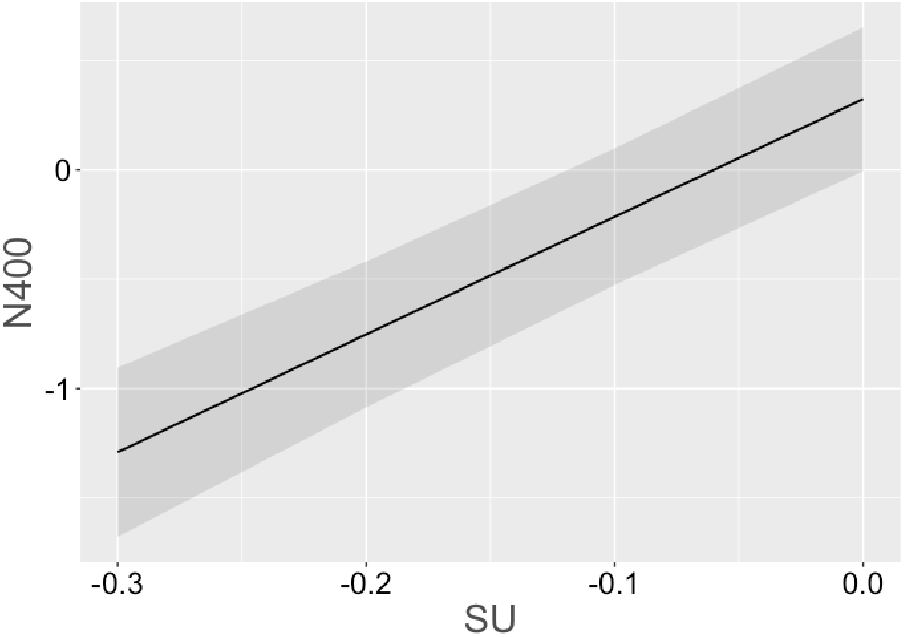
Relation between the amplitude of the N400 and the SU as estimated by the LME model described in Tab. 1.

In addition, we compared two separate models containing positional and stochastic measures, one predicting the amplitude of the N400, the other predicting the SU. The aim of this second analysis is to investigate whether the SU and the amplitude of the N400 are influenced in the same way by the stochastic and positional properties of the sentence. Both analyses are aimed at investigating the validity of the SU as approximation of the processing dynamics underpinning the generation of the ERP component under scrutiny.

### Predicting the N400

Tab. 1 contains the results of a linear mixed effect model (LME) predicting the N400 ERP component amplitude obtained from Frank et al. (2015) as a function of the SU over the stimulus words. SU is included together with the ERP baseline (the activity of the 100 ms leading to the onset of each word), which is not subtracted directly from the dependent variable, but instead included as a variable of no interest in the model. The model is fit with per-subject and per-word random intercepts.

**Tab. 1:** Results of a LME model fitted with the SU and aimed at predicting the amplitude of the N400 component.

| | $\beta$ | CI. | z | p |
| --- | --- | --- | --- | --- |
| ERPbase | -0.11 | -0.12 – -0.10 | -21.03 | <0.001 |
| **SU** | 0.07 | 0.05 – 0.08 | 9.52 | <0.001 |

The results in Tab. 1 clearly indicate that **SU** significantly predicts the amplitude of the N400 (β = 0.07, *z* = 9.52, *p* < 0.001). Larger word-wise updates of the SG layer representation correspond with stronger negative deviation of the ERP signal in the N400 time segment.

### Comparing the effect of surprisal and of SU as predictors of the N400

In order to assess the contribution of the SU on the amplitude of the N400 above and beyond the effect of surprisal, we fitted two nested linear mixed effects models, one (called *Null*) containing as predictors only surprisal, the other (*Full*) containing also SU. Both models were fit with random per-subject and per-word random intercepts. Table 2 reports the results of the log-likelihood test between the two models, showing the difference in model fit to be significant (χ^2^ = 79.03, *p* < .0001).

**Tab. 2:** Results of log-likelihood comparison between **Null** and **Full** model.

| model | AIC | BIC | loglik | $\chi^2$ | Pr |
| --- | --- | --- | --- | --- | --- |
| **Null** | 52354 | 51405 | -25671 |  |  |
| **Full** | 51277 | 51336 | -25631 | 79.03 | < .0001 |

Table 3 contains the β estimates for the *Full* model predictors. Even with the presence of surprisal (β = –0.06, *z* = –8.08, *p* < 0.001), SU makes a significant contribution to the amplitude of the N400 (β = 0.06, *z* = 8.90, *p* < 0.001).

**Tab. 3:** Results of a *Full* model fitted with both surprisal and SU and aimed at predicting the amplitude of the N400 component.

| | $\beta$ | CI. | z | p |
| --- | --- | --- | --- | --- |
| ERPbase | -0.11 | -0.12 – -0.10 | -21.03 | <0.001 |
| surprisal | -0.06 | -0.08 – -0.05 | -8.08 | <0.001 |
| **SU** | 0.06 | 0.05 – 0.08 | 8.90 | <0.001 |

### Comparing the effect of predictors on SU and N400

Tab. 4 and Fig. 6 contain the results of two linear models, one predicting SU, and the other predicting the N400 amplitude from Frank et al. (2015). As in the previous section, SU values were generated using the stimuli contained in Frank et al. (2015). Both models contain as predictors word position in the sentence, word frequency, and word surprisal. Word frequency was obtained from the BNC (Clear, 1993). Surprisal was computed using SRILM (Stolcke, 2002), also on the BNC. For the model predicting the N400 we also included the ERP baseline and per-subject and per-word random intercepts, as in the analyses reported in Tab. 1.

**Fig. 6:**
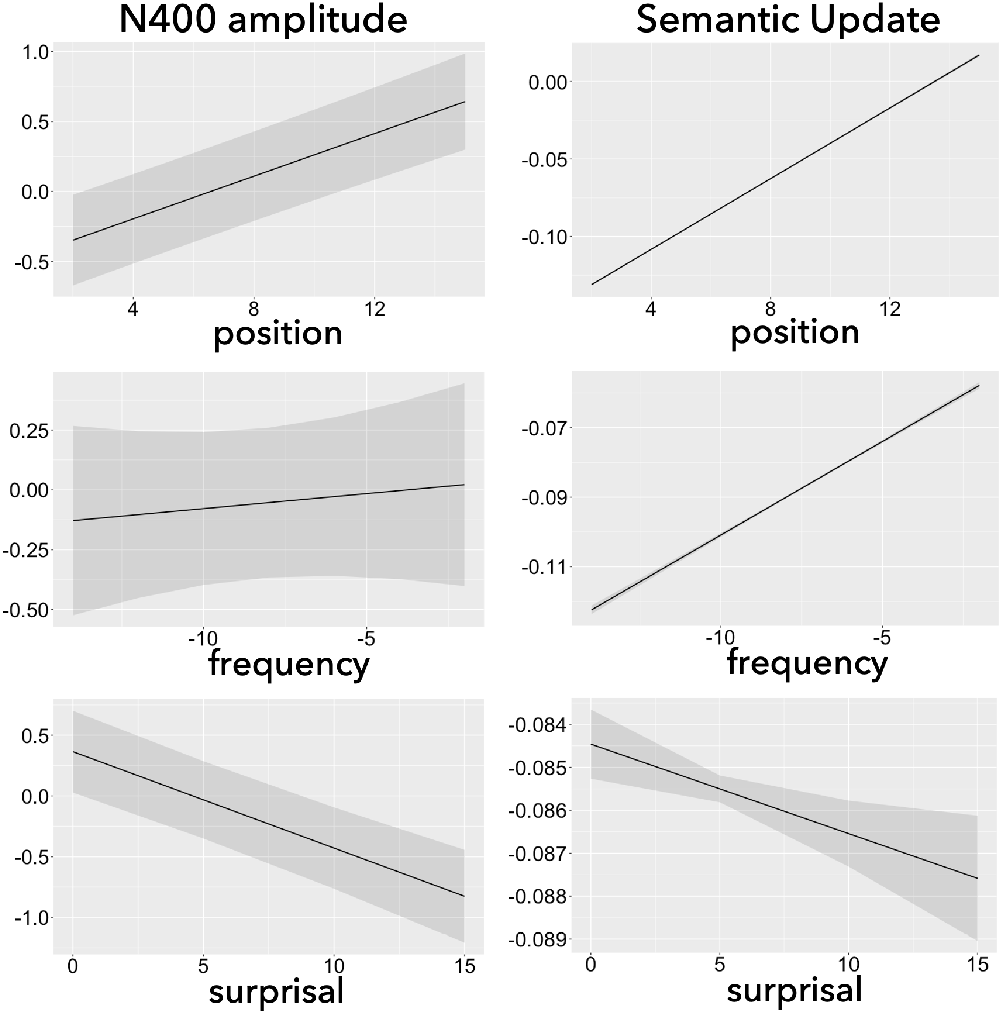
Influence of word position (1^*st*^ row), frequency (2^*nd*^ row), and surprisal (3^*rd*^ row) on the amplitude of the N400 (1^*st*^ column) and on the SU (2^*nd*^ column).

**Tab. 4:**
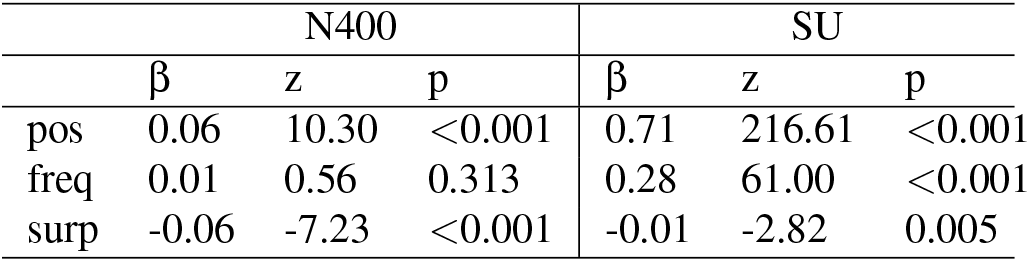
Influence of word position (pos), frequency (freq), and surprisal (surp) on the amplitude of the N400 and on the SU measure.

Word **position** in the sentence has a significant positive effect on both variables, indicating that the magnitude of these variables decreases as a function of the position of the word it is computed on (N400: β = 0.06, *z* = 10.30, *p* < 0.001, SU: β = 0.71, *z* = 216.61, *p* < 0.001). **Frequency** has a significant positive effect on SU, meaning that more frequent word forms elicit a smaller update of the SG layer representation (β = 0.28, *z* = 61.00, *p* < 0.001). Frequency shows also a positive, albeit not significant, effect on the amplitude of the N400 (β = 0.01, *z* = 0.56, *p* = 0.313). Please note however that other ERP studies did find significantly smaller N400 amplitudes for more high frequent words, which would be consistent with our modelling results (Rabovsky, Álvarez, Hohlfeld, & Sommer, 2008). **Surprisal** has a negative effect on the magnitude of the SU (β = −0.01, *z* = −2.82, *p* = 0.005), and on the amplitude of the N400 (β = −0.06, *z* = −7.23, *p* < 0.001). This indicates that the more unexpected a word is in a given context, the larger is the update of the inner representation generated by the SG model, and the larger is the negative deflection corresponding to the N400 evoked potential component. Overall, these results seems to indicate that both the SU and the N400 respond similarly to the effects of the position of a word in the sentence, its overall frequency and the probability of appearing in a specific context.

## Discussion

Previous studies have already suggested that N400 amplitudes reflect the change of a probabilistic representation of meaning corresponding to an implicit semantic prediction error. This was based on showing how the SU responds similarly as the N400 amplitude to a series of lexical semantic manipulations including semantic congruity, cloze probability, semantic and associative priming, and repetition, among others (e.g., Rabovsky et al., 2018). The analyses reported in this paper showed a significant relation between the amplitude of the N400 component and the update of the probabilistic semantic representation (SU) generated by a SG model trained on a large scale corpus of naturalistic texts (Tab. 1 and Fig. 5). Further analyses indicate that word position, word frequency and surprisal have, in relative terms, similar effects on the SU as they have on N400 amplitudes (Tab. 4 and Fig. 6). Both these analyses suggest that the SU is a valid approximate of the ERP component under examination. The fact that the update of a probabilistic semantic representation (the SU) observed in a corpus-trained SG model predicts the fluctuation of N400 amplitudes reinforces the intuition that the model itself is a valid approximation of sentence processing and that the N400 reflects an internal temporal difference prediction error at the level of meaning.

These results were obtained on electrophysiological data that was collected on quasi-naturalistic stimuli, i.e. stimuli that were not explicitly designed to elicit a strong N400 effect, but that were sampled in order to cover the natural complexity of linguistic material. This would not have been possible with the previous small scale implementation of the SG model. In general, despite advantages in terms of transparency and interpretability, one important limitation of small scale models trained on synthetic environments is the indirect relation between model and human data, making the testing of the hypothesis implemented in the model also somewhat indirect and based on the assumption that the small synthetic environment adequately captures the relevant statistical properties of human environments. A model trained on a large scale corpus allows to test the hypothesis implemented in the model in a much more direct way and thus seems crucial to rigorously test the model.

## References

Aurnhammer, C., & Frank, S. L. (2019). Comparing gated and simple recurrent neural network architectures as models of human sentence processing. In A. K. Goel, C. M. Seifert, & C. Freksa (Eds.), (pp. 112–118). Cognitive Science Society: Austin, TX.

Bentin, S., McCarthy, G., & Wood, C. C. (1985). Event-related potentials, lexical decision and semantic priming. Electroencephalography and Clinical Neurophysiology, 60(4), 343–355. doi: https://doi.org/10.1016/0013-4694(85)90008-2

Bojanowski, P., Grave, E., Joulin, A., & Mikolov, T. (2017). Enriching word vectors with subword information. Transactions of the Association for Computational Linguistics, 5, 135–146. doi: 10.1162/tacl_a_00051

Brouwer, H., Crocker, M. W., Venhuizen, N. J., & Hoeks, J. C. J. (2017). A neurocomputational model of the n400 and the p600 in language processing. Cognitive Science, 41, 1318–1352.

Brouwer, H., Delogu, F., Venhuizen, N. J., & Crocker, M. W. (2021). Neurobehavioral correlates of surprisal in language comprehension: A neurocomputational model. Frontiers in Psychology, 12.

Clear, J. H. (1993). The british national corpus. In The digital word: Text-based computing in the humanities (p. 163–187). Cambridge, MA, USA: MIT Press.

Collobert, R. (2011). Deep learning for efficient discriminative parsing. In International conference on artificial intelligence and statistics (AISTATS).

Collobert, R., Weston, J., Bottou, L., Karlen, M., Kavukcuoglu, K., & Kuksa, P. (2011). Natural language processing (almost) from scratch. Journal of Machine Learning Research (JMLR), 12, 2493–2537.

Fitz, H., & Chang, F. (2019). Language erps reflect learning through prediction error propagation. Cognitive Psychology, 111, 15–52.

Frank, S. L., Monsalve, I., Thompson, R., & Vigliocco, G. (2013). Reading time data for evaluating broad-coverage models of english sentence processing. Behavior research methods, 45, 1182—1190. doi: 10.3758/s13428-012-0313-y

Frank, S. L., Otten, L. J., Galli, G., & Vigliocco, G. (2015). The ERP response to the amount of information conveyed by words in sentences. Brain and Language, 140, 1–11. doi: 10.1016/j.bandl.2014.10.006

Hochreiter, S., & Schmidhuber, J. (1997, November). Long short-term memory. Neural Computation, 9(8), 1735–1780. Retrieved from https://doi.org/10.1162/neco.1997.9.8.1735 doi: 10.1162/neco.1997.9.8.1735

Kingma, D. P., & Ba, J. (2015). Adam: A method for stochastic optimization. CoRR, abs/1412.6980.

Kutas, M., & Federmeier, K. D. (2011). Thirty years and counting: finding meaning in the N400 component of the event-related brain potential (ERP). Annual review of psychology, 62, 621–47. doi: 10.1146/annurev.psych.093008.131123

Kutas, M., & Hillyard, S. A. (1984). Brain potentials during reading reflect word expectancy and semantic association. Nature, 307, 161–163.

McClelland, J. L., St. John, M. F., & Taraban, R. (1989). Sentence comprehension: A parallel distributed processing approach. Language and Cognitive Processes, 4, 287–335.

Merkx, D., & Frank, S. (2020). Comparing transformers and rnns on predicting human sentence processing data. ArXiv, abs/2005.09471.

Nivre, J. (2003). An efficient algorithm for projective dependency parsing. In Proceedings of the 8th international workshop on parsing technologies (IWPT 03) (pp. 149–160).

Palmer, M., Gildea, D., & Kingsbury, P. (2005). The Proposition Bank: An annotated corpus of semantic roles. Computational Linguistics, 31(1), 71–106. doi: 10.1162/0891201053630264

Parviz, M., Johnson, M., Johnson, B., & Brock, J. (2011). Using language models and latent semantic analysis to characterise the N400 neural response. In Proceedings of the australasian language technology association workshop 2011 (pp. 38–46). Canberra, Australia.

Rabovsky, M. (2020). Change in a probabilistic representation of meaning can account for N400 effects on articles: A neural network model. Neuropsychologia, 143, 107466. doi: https://doi.org/10.1016/j.neuropsychologia.2020.107466

Rabovsky, M., Hansen, S. S., & McClelland, J. L. (2018). Modelling the N400 brain potential as change in a probabilistic representation of meaning. Nature Human Behaviour, 2, 693–705.

Rabovsky, M., & McClelland, J. L. (2020). Quasi-compositional mapping from form to meaning: a neural network-based approach to capturing neural responses during human language comprehension. Philosophical Transactions of the Royal Society B: Biological Sciences, 375(1791), 20190313. doi: 10.1098/rstb.2019.0313

Rabovsky, M., Álvarez, C. J., Hohlfeld, A., & Sommer, W. (2008). Is lexical access autonomous? evidence from combining overlapping tasks with recording event-related brain potentials. Brain Research, 1222, 156–165. doi: https://doi.org/10.1016/j.brainres.2008.05.066

Sayeed, A., Shkadzko, P., & Demberg, V. (2018). Rollenwechsel-english: a large-scale semantic role corpus. European Language Resources Association. doi: http://dx.doi.org/10.22028/D291-30972

Stolcke, A. (2002). SRILM - an extensible language modeling toolkit. In INTERSPEECH 2002.

van Petten, C., & Kutas, M. (1990). Interactions between sentence context and word frequency in event-related potentials. Memory cognition, 18, 380–93. doi: 10.3758/BF03197127

